# Left Hippocampus to Anterior Cingulate Cortex Connectivity Correlates with Worse Recent Verbal Memory in Pornography Addicted Juveniles

**DOI:** 10.1101/2020.12.23.424123

**Authors:** Pukovisa Prawiroharjo, Rizki Edmi Edison, Hainah Ellydar, Peter Pratama, Sitti Evangeline Imelda Suaidy, Nya’ Zata Amani, Diavitri Carissima, Ghina Faradisa Hatta

**Affiliations:** Neurology Department, Division of Neurobehavior, Faculty of Medicine, Universitas Indonesia/Cipto Mangukusumo Hospital, Jakarta, Indonesia; Neuroscience Center, Universitas Muhammadiyah Prof. Dr. Hamka, Jakarta, Indonesia; Yayasan Kita dan Buah Hati, Bekasi, Indonesia; Independent scholar, Jakarta, Indonesia; Faculty of Psychology, Universitas Islam Negeri Syarif Hidayatullah, Jakarta, Indonesia

**Keywords:** Addiction, Juvenile, Pornography, Brain Connectivity, Cognitive Function

## Abstract

**Background and aims:** Imperil by the convenience of information and knowledge access, children exposed to pornography have worsened. As such, this study aims to gain insight into brain connectivity and cognitive function effect of pornography addiction in juveniles, as the best of our knowledge, this study is the first to specifically learn about memory function in juvenile’s pornography addiction.

**Methods:** We screened 30 juveniles with 4 dropouts (13 non-addiction vs 13 addiction group). Subjects underwent neuropsychiatric tests (memory, attention, and intelligence) and fMRI image acquisition. We carried correlation analysis of brain connectivity and neuropsychiatric test results.

**Results:** Significant disconnection between left hippocampus to ACC (Z-transformed r-value, non-addiction vs addiction = 0.07 ± 0.19 vs −0.08 ± 0.17, p=0.04, cohen d=0.83) followed by worse verbal recent memory in pornography addicted juveniles (RAVLT A6 sub-score, p < 0.01, d=0.67; A7 sub-score, p=0.01). Attention and intelligence test resulted to insignificant correlation.

**Discussion:** This data-driven analysis result strongly promotes the involvement of cortico-subcortical systems in pornography addiction, emphasizing the role of reward system pathology, indifferent to addiction pathophysiology in general. Decline in working memory, which are maintained by corticolimbic network, including hippocampus and ACC, affects goal-oriented behaviour greatly. This, correspond to our significant result of addiction group’s decline in memory, regardless of its association with attention and intelligence.

**Conclusion:** Disconnection between left hippocampus to ACC suggested similar neurobiological abnormalities as seen on other addictive disorders. Its disconnection was also correlated with worse verbal recent memory in pornography addicted juveniles, without affecting attention and intelligence, results showed.

## INTRODUCTION

Era of globalization, which eases access to information and knowledge through the Internet, has unfortunately exposed our children to pornography, intentionally or unintentionally. Unintentional exposure may risks from mistyping website addresses, searching for terms with or without sexual meaning, or accidentally encountering pop-up images and advertisements [1]. In virtue of this matter, it is important to consider developmental factors in cognitive processing, as children aged less than 7 years old, have difficulty in differentiating between the on-screen and real-life situation. Incomplete cognitive development of children and adolescents may pose real risks in affecting how pornography itself is perceived and acted upon by children in their daily life, thus in turn may develop into problematic behaviours [2]. Problematic sexual behaviour in young children aged under 12 years old are thought to be the results of several factors, of which is pornography viewing, a study showed [3].

Studies in U.S. showed pornography exposure in children varies between studies, with unintentional exposure ranged from 19-34% [4 – 6], and intentional exposure 43% [4]. Indonesia is no exception, in 2012, at least 50% Indonesian juveniles aged 10-19 y.o. has viewed pornography contents, 14% intentionally exposed. This became worse in 2016, where at least 97% elementary students of 4–6^th^ grade in Jakarta and its surrounding had been exposed to pornography [7].

Since long ago, substance addiction has gathered significant interests of both researchers and social activists. However, there are still very few studies concerning pornography addiction, especially in the field of neuroscience. Addiction are thought to be affiliated with the reward system of the brain, which involves emotion and executive functions of the brain--the amygdala, hippocampus, and frontal cortex [8, 9]. Studies have found that dysfunction in prefrontal cortex (PFC), a region of brain, is accounted for the reduction in executive functions and behavioural inhibitory control [10 – 12]. Despite the lack of neuroscientific studies of pornography addiction, even more studies focusing on the younger population, a growing body of evidence suggests that the mechanism behind pornography addiction is indifferent of substance addiction, i.e. both involves disruption in PFC area [13 – 15]. The involvement of both emotion and executive functions in neurobiology of addiction are thought to be connected through the broader form of corticolimbic network, in which are anterior cingulate cortex (ACC), hippocampus, and prefrontal cortex (PFC) that may intercorrelate with each other [8, 16, 17].

As those affected areas turns into decline, it may affect its function, in this matter are the children’s and adolescents’ who have still substantial room of growth in life, therefore may lead them to life with cognitive or behavioural problems. Studies also found that pornography addiction is among the causes of social problems [18,19]. The lack of scientific studies addressing these arising problem of the youths concerns us. Therefore, this study aimed to use functional MRI in investigating brain connectivity, especially in the area of PFC, ACC, and hippocampus, which thought to be much affected; and its possible disruption of affiliated cognitive function— memory, intelligence, and attention---in pornography addicted juveniles.

## MATERIALS AND METHODS

### Participants

We used the data from 30 juveniles aged 12-16 years old from our previous study, recruited during December 2017-February 2018, in various events held by YKBH in Bekasi, Indonesia. Subjects were grouped into pornography addiction and non-addiction group, which were determined using Pornography Addiction Test, a battery of neuropsychological test designed and validated by YKBH [20]. Exclusion criteria were left-handed, incomplete tests/fMRI procedures, verbal or language disorder, history of brain-related disorder or disease, head trauma, trauma during pregnancy or birth, developmental, psychological or neurological disorder, or mental illness.

### Procedures

All participants underwent pornography addiction screening, memory function assessment by Ray Auditory Verbal Learning Test (RAVLT) for auditory-verbal memory and Ray-Osterrieth Complex Figure Test (ROCFT) for visual memory, Wechsler Intelligence Scale for Children for IQ (WISC IQ), and fMRI image acquisition. Each subject underwent MRI scanning using a 3-Tesla scanner (GE Advance Workstation 4.5). We first performed an initial scan to center the field of view on the subject’s brain. We then performed a 3D T1-weighted turbo field echo scan for anatomical reference (repetition time [TR] = 8.3 ms; echo time [TE] = 3.2 ms; FOV = 240 mm; matrix size = 256 × 256; slice thickness = 3.0 mm, space 1 mm). Lastly, we performed a gradient echo-planar sequence for functional imaging (TR = 3000 ms; TE = 30 ms; flip angle = 90 degrees; FOV = 240 mm; 36 x 3 mm slices; space 1mm) which lasted for 7 minutes. Subjects were asked to refrain from any psychoactive substance use and sexual activity during the 24 hours preceding fMRI.

### Measures

#### Pornography Addiction Screening

We completed the pornography addiction screening with a self-reported questionnaire, which developed by expert psychologist. Several indicators commonly found in juveniles with high pornography consumptions were emphasized, based on field studies and literature researches. Those are grouped into three dimensions: 1) time spent on pornography in the last six months, consisted of number of times, frequency, and duration spent; 2) motivation to use pornography, consisted of factors encouraging access to pornography, which includes sexual curiosity, emotional avoidance, sensation seeking, and sexual pleasure; and 3) problematic pornography use, consisted of functional and distress problems, excessive use, control difficulties, and using to escape negative emotions. The questionnaire consisted of 92 items and has been tested on 740 grade six to ten students in Indonesia, which further details corresponded in an unpublished report **(Table S1 and S2).**Three additional questions were added to minimize faking good possibility, thus we excluded subjects who answered these according to social desire. Pornography addiction was defined as those with weighted score greater than or equal to 32. Psychometric analysis exhibited valid (CFA > 1.96) and reliable (Cronbach’s Alpha > 0.7) for all items in the questionnaire.

Being specially developed and adapted to juveniles population, the questionnaire was very suitable for this pornography addiction study. Furthermore, the additional questions to exclude subjects who faked good and forced choice technique questions in this questionnaire allowed for less bias. The limitation of this questionnaire may be its number of questions, as it may induce fatigue and boredom on the subjects. Although it being specially adapted to juveniles population made it very suitable for this study, its use in other context outside of juvenile pornography addiction may require wording adjustment to make a better understanding of the vocabularies used.

#### Memory Assessments

All subjects underwent memory function assessment by Ray Auditory Verbal Learning Test (RAVLT) for auditory-verbal memory and Ray-Osterrieth Complex Figure Test (ROCFT) for visual memory. All subjects were first instructed to copy the figure in ROCFT sheet onto a blank paper. After the direct copy, subjects continued to undertake RAVLT. With a presentation rate of one word per second, the first 15 noun-word list (Indonesian version) was read to participants (list A). Subsequently the presentation, subjects were requested to recall as many words as possible, regardless the word order. The steps were repeated 5 times, with each trial recall was recorded. Then, the next list (list B) was presented, subjects were also requested to recall as many words as possible. Promptly after recalling list B, the subjects were again requested to recall list A, twice, with 30 minutes interval in between (short recall, A6; long recall, A7). Finishing the RAVLT, which would take approximately 30-40 minutes to complete, all subjects were requested to reproduce the figure from memory of previous ROCFT. Attention, as a factor influencing working memory, [21,22] was evaluated using Trail Making Test (TMT) A and B. The subjects were instructed to connect a set of 25 dots of numbers (TMT A) and mix of numbers and letters, alternately (TMT B), after finishing each 8-dot-set-sample. The time taken of each set was recorded.

#### Intelligence Measurements

All subjects’ intelligence were measured by Wechsler Intelligence Scale for Children for IQ (WISC IQ) that was designed for children between 5 and 16 years of age. The test consists of 10 basic tests and 2 supplementary tests. Basic tests comprise of verbal tests (information, vocabulary, arithmetic, comprehension and similarities) and performance tests (picture completion, picture arrangement, block design, object assembly, and digit symbol). Supplementary tests comprise of (verbal tests, digit span, performance test and maze).

### Analysis

#### Imaging Analysis

Data pre-processing followed a standard pipeline, consisting of realignment and unwarping, slice-timing correction, normalization, outlier detection, and finally spatially smoothing (full width at half maximum = 10 mm) which resulted in both functional and structural images in MNI-space. All were conducted using CONN toolbox version 17.f (https://www.nitrc.org/projects/conn) in SPM version 12 (http://www.fil.ion.ucl.ac.uk/spm/). De-noising which consisted of removing white matter and cerebrospinal fluid noise with 5 dimensions each, scrubbing, motion regression, band-pass filtering (0.01-0.10 Hz), and linear detrending was later performed. Nine nuisance covariates (time-series predictors for global signal, white matter, cerebrospinal fluid, and the six movement parameters) were sequentially regressed from the time-series.

Previous studies have shown patients with pornography addiction have impaired control and impulsivity, much associated with subcortical structures connectivity, such as hippocampus, putamen, and anterior cingulate cortex, thus were chosen as the seed regions and targets in ROI-to-ROI analysis (**Fig 1**).[8, 13, 23-26] Correlation map was produced by computing the Pearson linear correlation coefficients between the time course of signal in each ROI and the average signal of the seed. The process was calculated using CONN. To elucidate significant connection in each ROI-to-ROI group, one-sample T-Test were used. The ROI analysis would include the targets in cases where it was revealed significant connections between target and seed regions, which were defined anatomically from the MNI template.

**Fig 1.**
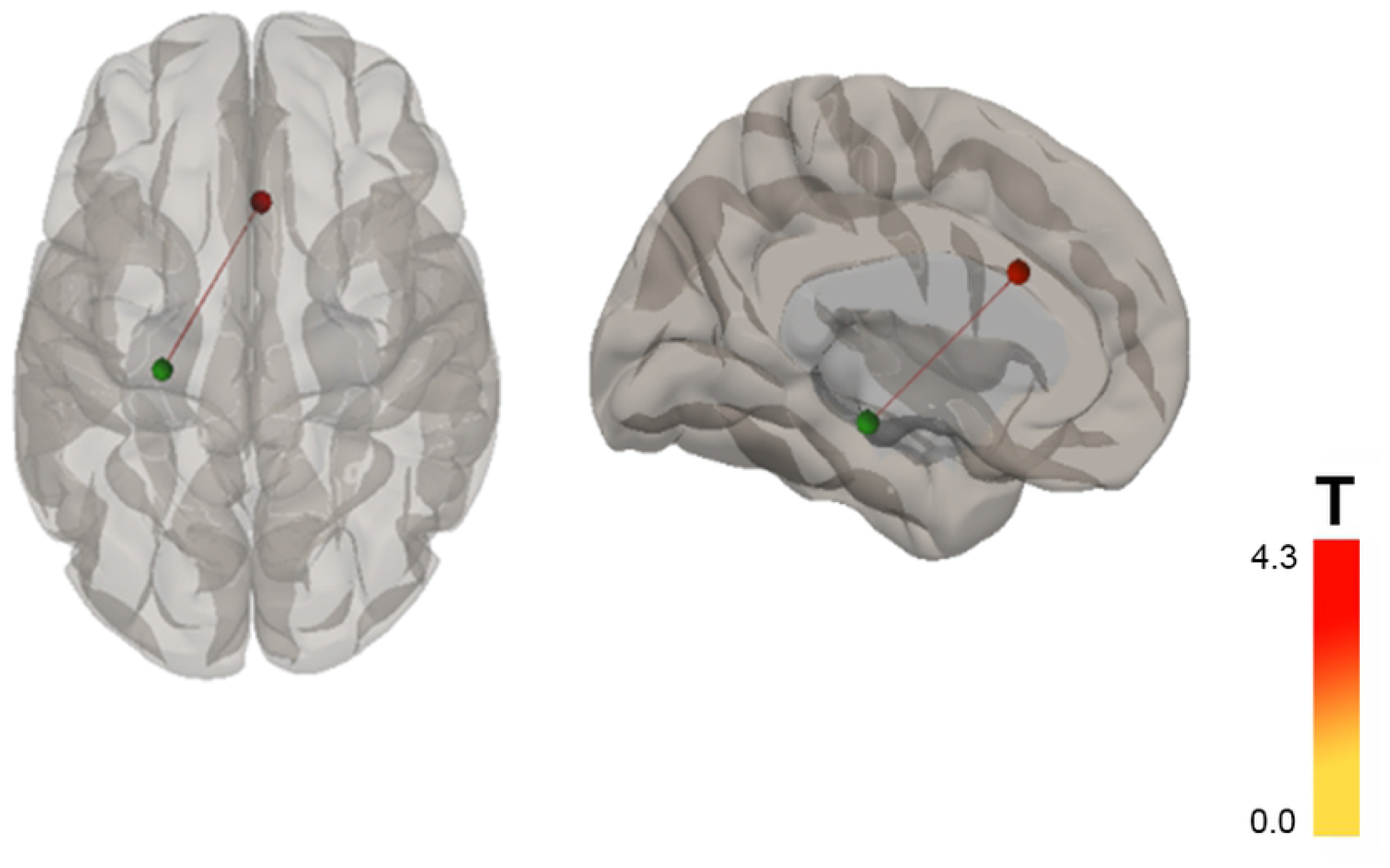
ROI seed (left hippocampus, green) and target (ACC, red), from (a) superior view, and (b) left medial view.

To improve the normality of the distribution, we converted the correlation coefficients to Fisher’s Z-transformed r-values. We then performed correlation analysis between the resulting Z-transformed r-values of significant ROI-to-ROI analysis (non-addiction vs addiction group) with the significant neuropsychiatric test and each domain sub-scores. For that purpose, we performed Spearman rank test.

#### Statistical Analysis

We used Mann-Whitney tests for comparison between groups. Correlation analysis was performed using Spearman rank test. Statistical significance was assumed on p-value of < 0.05. Cohen d’s effect size was also calculated for each result, assuming d-value of >0.2 as small, >0.5 as medium, and >0.8 as large effect size. All statistical analysis was performed SPSS version 21.

### Ethical Approval

The study was approved by Health Research Ethical Committee of Faculty of Medicine Universitas Indonesia (Clearance No. 1155/UN2.F1/ETIK/2017) and conducted in accordance to Helsinki Declaration. No subject was confronted with pornographic material in this study. Informed consent was obtained from all participants, represented by respective parents.

## Results

### Demographic Data

From the initial 30 subjects, 4 was dropped due to incomplete fMRI data, resulting in final 26 subjects (13 non-addiction vs 13 addiction group). Demographic and neuropsychiatric test results were shown in **Table 1**. Age was matched between both groups (p = 0.34). Among memory test results, there was significant difference in RAVLT A6 subscore (non-addiction vs addiction, A6 = 14.08 ± 1.12 vs 11.15 ± 2.19, p < 0.01, d = 1.68) and A7 subscore (A7 = 14.23 ± 0.93 vs 11.69 ± 2.63, p = 0.01, d = 1.28). There was no significant difference between groups in other scores of RAVLT, ROCFT, TMT A, TMT B, and WISC IQ results.

**Table 1.**
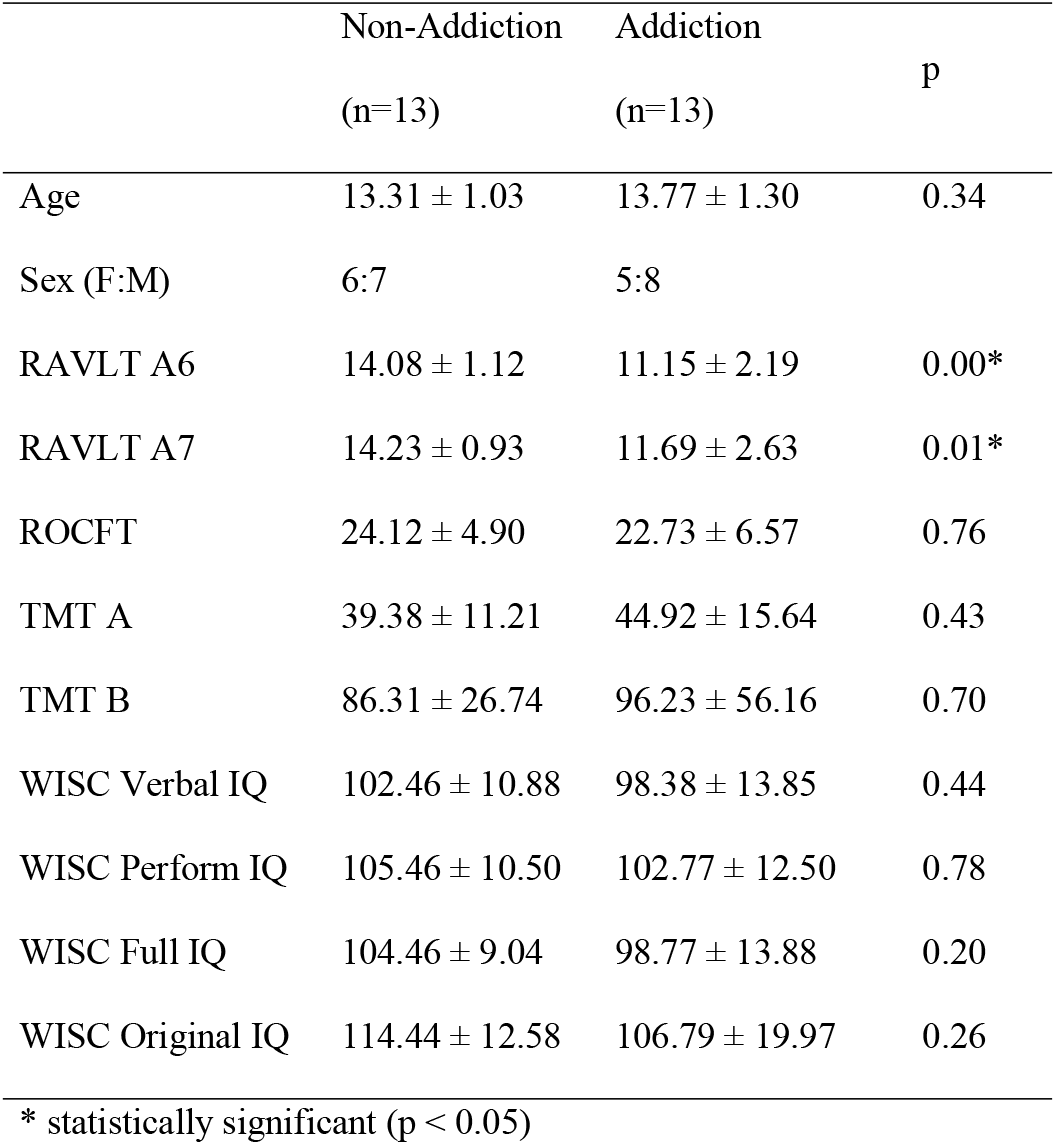
Demographic and neuropsychiatric tests comparison.

### Brain Connectivity

From ROI-to-ROI analysis in CONN with non-addiction>addiction contrast, we found one significant connection difference, which was left hippocampus to ACC (Z-transformed r-value, non-addiction vs addiction = 0.07 ± 0.19 vs −0.08 ± 0.17, p = 0.04, d = 0.83) (**Fig 2**). Thence, it showed hypoconnectivity between hippocampus to ACC in addiction group.

**Fig 2.**
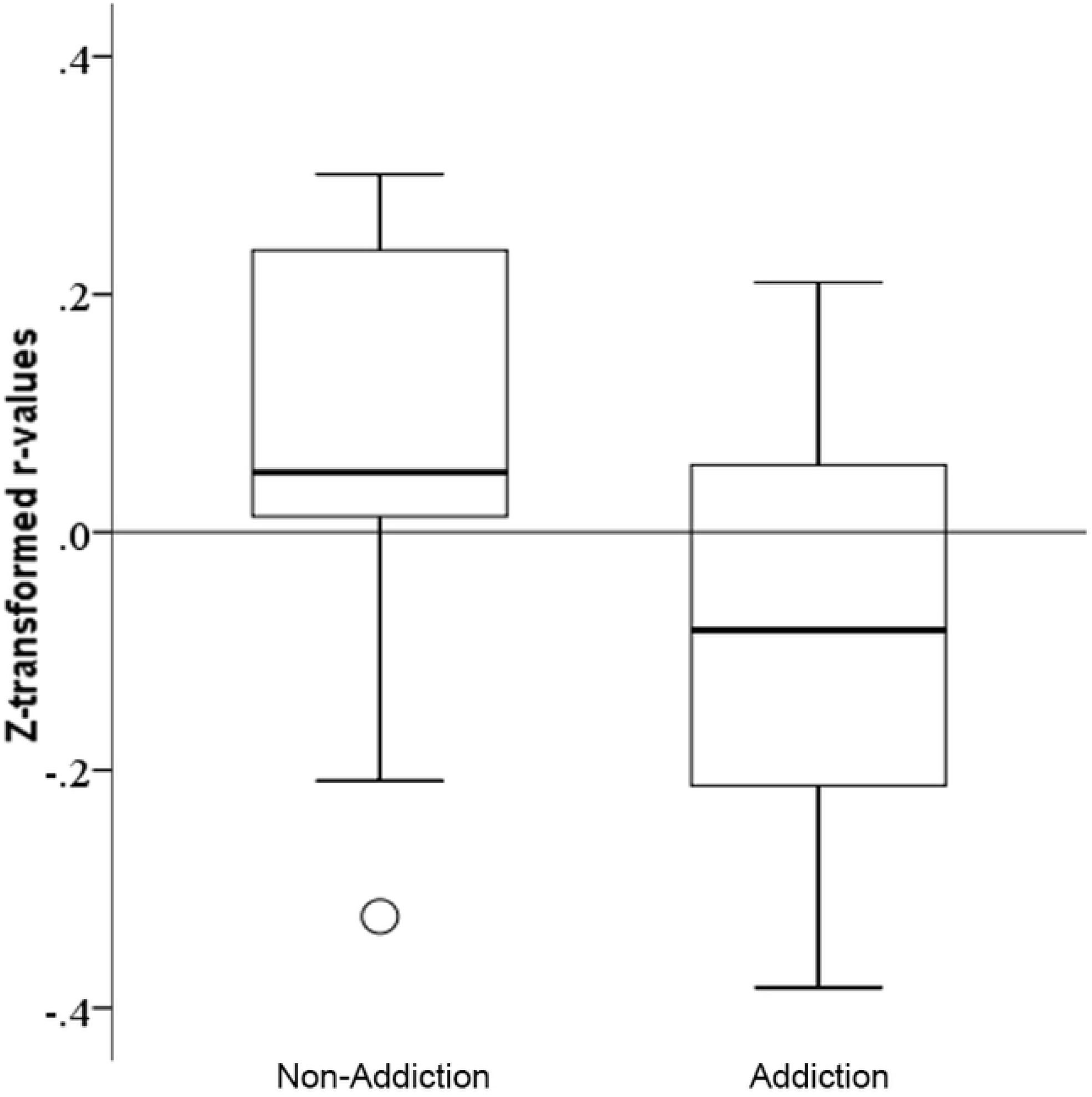
Boxplot comparison of Left Hippocampus-ACC Z transformed r-values connectivity between Non-Addiction and Addiction groups.

### Correlation Analysis

We performed Spearman rank test between significant neuropsychiatric test (RAVLT A6 and A7 score) and Z-transformed r-value of significant connectivity (left hippocampus to ACC) in pornography addiction group, and found significant correlation with A6 (r = 0.43, p = 0.04, d = 0.67) but not A7 (p = 0.10) (**Fig 3**).

**Figure 3.**
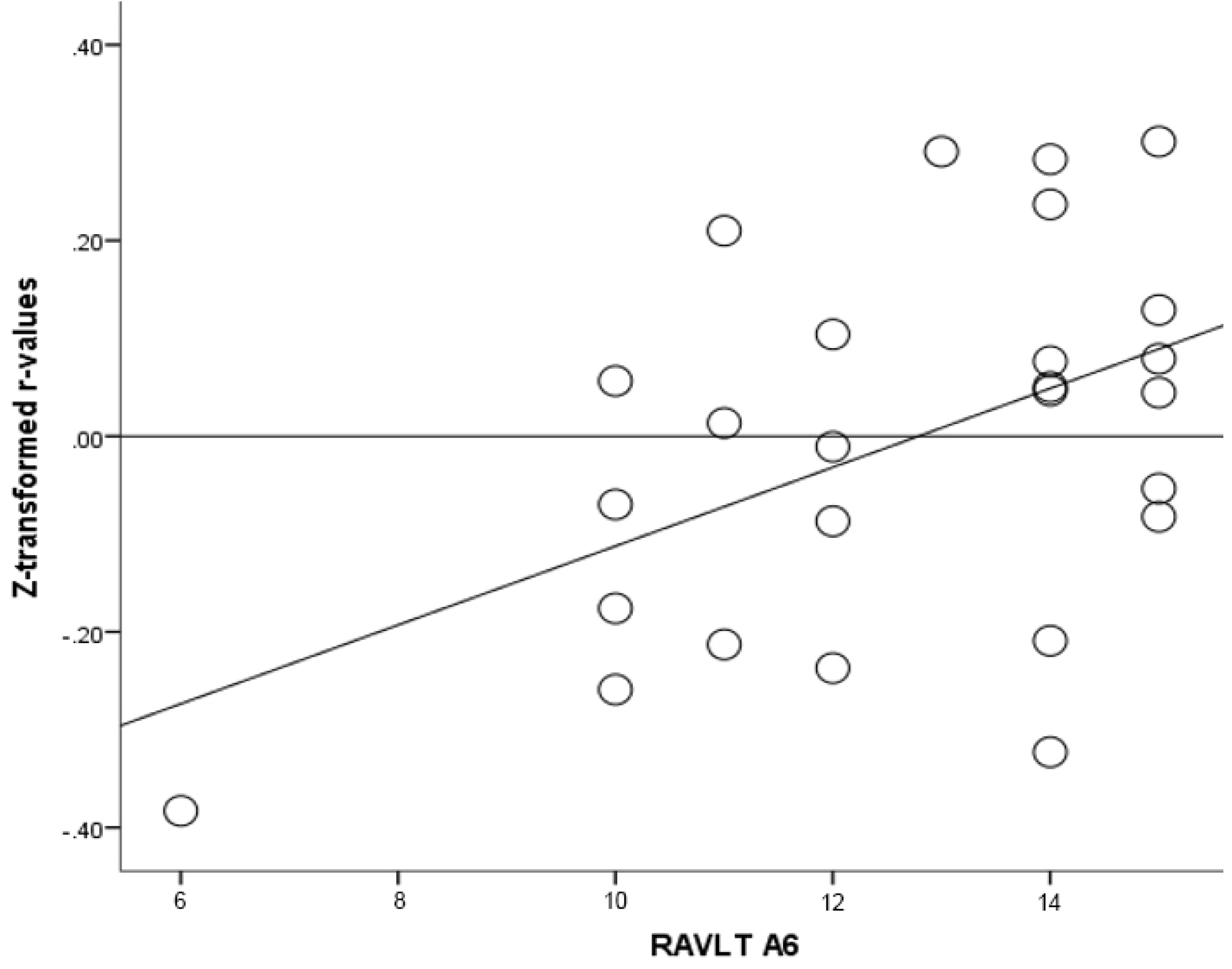
Scatterplot of Left Hippocampus-ACC Z transformed r-values connectivity with A6 score of RAVLT.

## Discussion

Addiction is explained by American Society of Addiction Medicine (ASAM) as a primary, chronic disease of brain, related to reward, motivation, and memory circuit. Dysfunction of such circuit leads to characteristic biological, psychological, social and spiritual manifestations—thus reflected in an individual pathologically seeking reward and/or relief by substance use or other behaviours.[27]

In terms of addiction pathophysiology, emotion and cognitive function have long been ascribed taking part within. Mesolimbic dopamine pathway is thought affected the neurobiology of addiction. The pathway particularly connects with three other key regions to form integrated circuits, commonly called as the reward system: the amygdala (emotions, includes positive and negative, also emotional memory), hippocampus (long term memories processing and retrieval), and the frontal cortex (behaviour coordination). As aforementioned above, mesolimbic dopamine pathway takes a role in addiction and reward system. A study showed continued release of dopamine into the reward system in individual compulsively watching internet pornography that it promotes neuroplastic changes reinforcing the experience. Excessive behaviours influenced neuroanatomical and reward system changes [8, 9]. Taking part in the reward system, hippocampus is important for long-term memory, contextual, spatial, and emotional processing; whilst the executive function aspect was taken in by the frontal cortex [8, 16, 28-30]. PFC is particularly the key for aspects of working memory, temporal processing, decision making, flexibility and goal-oriented behaviour. As connectivity is a key determinant to functional intercourse within the brain, hippocampus-PFC’s role has been ascribed in the broader form of corticolimbic network, in which anterior cingulate cortex (ACC), hippocampus, and PFC may intercorrelate with each other.[8, 16, 17] The cingulate cortex is divided into four functionally distinctive regions, which comprises of Broadmann’s areas 24, 25, 32, and 33. It consists of the ACC, midcingulate cortex (MCC) or dorsal anterior cingulate cortex (dACC), posterior cingulate cortex (PCC), and retrosplenial cortex. Being a part of corticolimbic network, the cingulate cortex has role both in emotional and cognitive function. ACC and dACC in particular is thought to carry out reward-based decision making. It has been shown that ACC receives projections from structures that process rewards, i.e. striatum, mesolimbic dopamine system, dorsolateral prefrontal cortex, orbitofrontal cortex, and the supplementary and primary motor cortices [31, 32].

Various evidence supports the interaction of hippocampus-PFC in reward learning, thus intercorrelate with addiction and impulsivity [17,33]. Particular region in hippocampus has been implicated in context-dependent process, which is the ventral subiculum of the hippocampus. Studies have shown its inactivation decreases reinstatement, thus marking its important role in drug-seeking behaviour [33]. Similar process has been shown in other kinds of addiction, i.e. game, internet, and pornography addiction [8, 12, 17, 26, 34-36]. Decline in working memory affects goal-oriented behaviour greatly, as aforementioned maintained by PFC and corticolimbic network [8, 16], thus suggested the problem existence in pornography addiction [37].

As shown in both internet and drug addiction, studies showed a clear link between addiction and aberrant connectivity in response of inhibition network, resulting in behavioural disorder that fail to inhibit unwanted actions, thus associated with poor impulse control. Studies have revealed lower grey matter density, abnormal white matter fractional anisotropy, reduced orbitofrontal cortical thickness, impaired brain activity, and nonetheless decreased functional connectivity.[8, 38] A study showed rather increased functional connectivity in right hippocampus, and decreased functional connectivity in right dACC and left caudate in the DMN. This abnormal functional organization of the DMN are discussed as addiction-related increased memory processing with decreased cognitive control—thus once again implying lack of impulse control, self-monitoring, and attention [36].

Of various studies about addiction, many showed consistent findings in which decreased functional connectivity in reward processing circuit, nonetheless between hippocampus and ACC. Although drug addiction studies are appreciably more dominating than other kinds of addiction, similar findings have also been shown on other addictions, i.e. game, internet, and pornography addiction [8, 12, 17, 26, 34-36]. The first fMRI study focusing on internet pornography addiction was published in 2014 and showed the same brain activity as seen in substance addicts and alcoholics [23].

Evidence of decreased brain functional connectivity is consistent with current models emphasizing the role of reward system pathology in addiction. This data-driven analysis result strongly promotes the involvement of cortico-subcortical systems in pornography addiction as the fact it emerged as the prominent pathology. Thus, indicating similar neurobiological characteristics with other addictive disorders, as such subcortical regions may play the core role of brain network pathology. Our study specifically enlisted juveniles as our focus of research, thus, being the first to specifically learn about memory function in juvenile’s pornography addiction, we were unable to directly compare our results to previous studies. Thence, we attempted to compare our results with other related studies. Our correlation analysis showed significant results in RAVLT A6 and A7 sub-score. On the other hand, ROCFT, which was to assess visual memory, showed no significant results. Attention, considered as a factor influencing memory, also left with no significant results, as seen on TMT A and B results. Attention ensures the selection of information we received, and later selected to be brought into working memory from perception or long term memory. That is to say, the two compounds do affect each other [39]. Therefore, our results exhibited significant decline in memory in addiction group, regardless its association with attention considered. In addition, another cognitive test that was assessed was WISC IQ test, also left no significant result. Therefore, it showed that addiction affected visual-auditory memory of cognitive function without affecting intelligence.

To hold up our results, previous studies had also found lower working memory in substance addiction [40-42], but not pathological gambling [40, 41]. Study by Nie, et al also showing similar decline in verbal memory of internet addiction group [43]. To complement these results, we did correlation analysis and found significant correlation of left hippocampus to ACC hypoconnectivity to RAVLT A6 sub-score decline (r = 0.43, p = 0.04, d = 0.67). Similar results of unaffected intelligence in internet addiction and non-addition group were also found by Park, et al, also tested with WISC IQ test with participants’ mean age of 15.17 years old [44]. Considering its pathology similarities with pornography addiction [8, 23], their similar results may as well be regarded.

Our enrolment of juveniles participants serves both as our study’s limitation and strength. As our population size (n = 13 of each group) was rather small, to avoid lacking in substantive significance, we also calculated the Cohen d’s effect size of each result. The effect size calculations show that the sample size of our study is appropriate. Being the pioneer study of pornography addiction in juveniles, we aimed to serve information for the most critical phase, in which are still growing and developing juveniles’ brains, thus might compensate underlying brain impairment.

## Conclusion

Our results suggest that adolescents with pornography addiction exhibit disconnection between left hippocampus to ACC that greatly play a role in reward system. This is consistent with other studies showing similar neurobiological abnormalities between pornography addiction and other addictive disorders. Given to its role in intelligence, memory, and learning, our results suggested that pornography addiction in juveniles correlates with worse verbal recent memory. Although, we did not find any significant correlation with intelligence and attention. Further research with larger population might aid to more defined results.

## Supporting Information

**Table S1. Mean performance on each of questionnaire’s components and concurrent validity, as opposed to the three-step-norm.** The 92-item questionnaire was followed by interview as qualitative input as well as questionnaire about children tendencies in sexual activities (**Table S2**) to confirm the screening (three-step-norm). Confirmatory factor analysis (CFA > 1.96) and Cronbach’s Alpha (0.903; CA > 0.7) values showed its validity and reliability.

**Table S2. Children tendencies in sexual activities questionnaire.** It ranged from never to very frequent in a scale of 1 to 5. Minimum score of 11 and maximum score of 55.

## Acknowledgements

The authors would like to thank Alexandra Chessa, Resti Siti Saleha, Kevin Widjaja, and Nia Soewardi for their contributions in this paper.

## Authors’ contribution

Conceptualization: PuP, REE, HE. Investigation: PuP, REE, HE, SEIS, NZA, DC.

Methodology: PuP. Formal analysis: PeP, GFH. Resources: HE, SEIS, NZA, DC.

Writing (draft preparation, review, and editing): PeP, GFH, PuP. All authors critically reviewed and approved the final version of the manuscript.

## ABBREVIATION LISTS

ACC: Anterior Cingulate Cortex
ASAM: American Society of Addiction Medicine
dACC: Dorsal Anterior Cingulate Cortex
DMN: Default Mode Network
fMRI: Functional Magnetic Resonance Imaging
FOV: Field of View
ICA: Independent Component Analysis
MCC: Midcingulate Cortex
MNI: Montreal Neurological Institute
NFC: Negative Functional Connectivity
PCC: Posterior Cingulate Cortex
PFC: Prefrontal Cortex
RAVLT: Ray Auditory Verbal Learning Test
ROCFT: Ray-Osterrieth Complex Figure Test
ROI: Region of Interest
TE: Echo Time
TMT: Trail Making Test
TR: Repetition Time
WISC IQ: Wechsler Intelligence Scale for Children for IQ
YKBH: Yayasan Kita Dan Buah Hati

